# Site-specific transgenesis of the *D. melanogaster* Y-chromosome using CRISPR/Cas9

**DOI:** 10.1101/310318

**Authors:** Anna Buchman, Omar S. Akbari

**Affiliations:** Section of Cell and Developmental Biology, University of California, San Diego, La Jolla, California, 92093, United States of America.; Tata Institute for Genetics and Society, University of California, San Diego, La Jolla, CA 92093, United States of America.

**Keywords:** RISPR, Cas9, Y-Chromosome, Drosophila, transgenesis, HDR

## Abstract

Despite the importance of Y-chromosomes in evolution and sex determination, their heterochromatic, repeat-rich nature makes them difficult to sequence and genetically manipulate, and therefore they generally remain poorly understood. For example, the *D. melanogaster* Y-chromosome, one of the best understood, is widely heterochromatic and composed mainly of highly repetitive sequences, with only a handful of expressed genes scattered throughout its length. Efforts to insert transgenes on this chromosome have thus far relied on either random insertion of transposons (sometimes harboring ‘landing sites’ for subsequent integrations) with limited success or on chromosomal translocations, thereby limiting the types of Y-chromosome related questions that could be explored. Here we describe a versatile approach to site-specifically insert transgenes on the Y-chromosome in *D. melanogaster* via CRISPR/Cas9-mediated HDR. We demonstrate the ability to insert, and detect expression from, fluorescently marked transgenic transgenes at two specific locations on the Y-chromosome, and we utilize these marked Y-chromosomes to detect and quantify rare chromosomal nondisjunction effects. Finally, we discuss how this Y-docking technique could be adapted to other insects to aid in the development of genetic control technologies for the management of insect disease vectors and pests.

## Introduction

The ability to perform site-specific transgenesis in insects and other organisms has greatly improved researchers’ ability to conduct controlled experiments in a precise way. In *D. melanogaster*, for example, prior to the advent of site-specific transgene integration technologies, most transgenesis was carried out using transposon-based vectors that inserted embedded transgenes randomly throughout the genome, leaving them vulnerable to distinct position effects arising from surrounding *cis*-regulatory regions and chromatin structure (Wimmer 2005). The development of site-specific transgenesis techniques such as those that rely on the phage φC31–derived site-specific integrase (Groth et al. 2004) or on recombinase-mediated cassette exchange (RMCE, (Oberstein et al. 2005)), represented a major advance (Wimmer 2005; Venken & Bellen 2005). This is because these techniques enabled more accurate and detailed transgene comparison and identification of insertion loci that are conducive to robust transgene expression levels and are fitness neutral, factors that can be especially important in practical applications of insect transgenesis such as vector control (Irvin et al. 2004; Wimmer 2005). However, the widely adopted site-specific recombinase-based transgenesis techniques (e.g., φC31) are limited by their requirement for pre-existing ‘landing sites’ within genomes of interest, which must first be generated by random transposon-mediated transgenesis, and unfortunatly these sites are lacking in most non-model organisms (Wimmer 2005).

The arrival of CRISPR technologies heralded a new era not only for traditional genome manipulation such as mutagenesis, but also for precise, site-specific transgenesis (Gratz et al. 2015; Bier et al. 2018). By using a simplified two-component system consisting of a *S. pyogenes* Cas9 endonuclease (SpCas9) and a single chimeric guide RNA (Jinek et al. 2012), one can generate DNA double-strand breaks (DSB) in a location of one’s choosing, provided it contains a 3-bp (NGG) protospacer adjacent motif (PAM) required for CRISPR/Cas9 function (Jinek et al. 2012). Then, as long as a donor template comprising homology-containing stretches (i.e., homology arms) flanking the target transgene is provided, the DSB break can be repaired via the cell’s own homology-directed repair (HDR) pathway to integrate the transgene at the precise location of the DSB (Gratz et al. 2015). This technique has been shown to be highly efficient in *D. melanogaster* (Gokcezade et al. 2014; Gratz et al. 2014; Gratz et al. 2015), and given CRISPR’s functionality in many insects (Gantz et al. 2015; Li, Bui, Yang, et al. 2017; Hammond et al. 2016; Dong et al. 2018; Sun et al. 2017; Li, Au, et al. 2017; Kohno et al. 2016; Li, Bui & Akbari 2017; Li et al. 2018), should be broadly applicable for insect transgenesis in general (e.g., (Li, Bui, Yang, et al. 2017; Hammond et al. 2016; Gantz et al. 2015).

The Y-chromosome of *D. melanogaster*, and of most other heteromorphic sex chromosome-bearing organisms, has so far presented an elusive target for site-specific transgenesis applications. The Y-chromosome’s degenerative, repetitive nature renders it difficult to sequence, assemble, and analyze genomically (Carvalho 2002; Piergentili 2010; Hall et al. 2016); additionally, it is almost entirely heterochromatic (Elgin & Reuter 2013), which likely makes it refractory to transgene integration and/or robust transgene expression (Bernardini et al. 2014). These factors have made it difficult to manipulate the Y-chromosome in a precise fashion, and the only successful attempts at placing transgenes on the Y-chromosome in insects, for example, have relied on random (and rare) events, such as chromosomal rearrangements or low frequency Y-specific transposon-mediated integrations, and subsequent manipulations (Starz-Gaiano et al. 2001; Szabad et al. 2012; Bernardini et al. 2014; Bernardini et al. 2017; Zhang & Stankiewicz 1998; Zhang & Spradling 1994).

Despite the challenges presented by the nature of Y-chromosome, having the ability to engineer it in *D. melanogaster* and other insects would be of utility in a number of different applications. Firstly, the ability to mark the Y-chromosome with an easily-scorable, specific marker may in itself be useful. For example, it would allow for the facile identification of rare Y-bearing or Y-lacking individuals (e.g., XXY females and XO males arising from nondisjunction, Figure 2A; Bridges 1913; Bridges 1916), which can be of experimental use (e.g., Brown & Bachtrog 2017; Lemos et al. 2010; Hearn et al. 1991; Yang et al. 2012). A marked Y-chromosome would also enable interspecific Y-chromosome introgression studies, which can shed light on Y-chromosome evolution and function (Araripe et al. 2016; Sackton et al. 2011; Bernardini et al. 2017), and can also be useful for sexing embryos (Bernardini et al. 2014; Condon et al. 2007), although it would also mark XXY females and fail to identify XO males, which may be problematic for certain applications.

**Figure 1.**
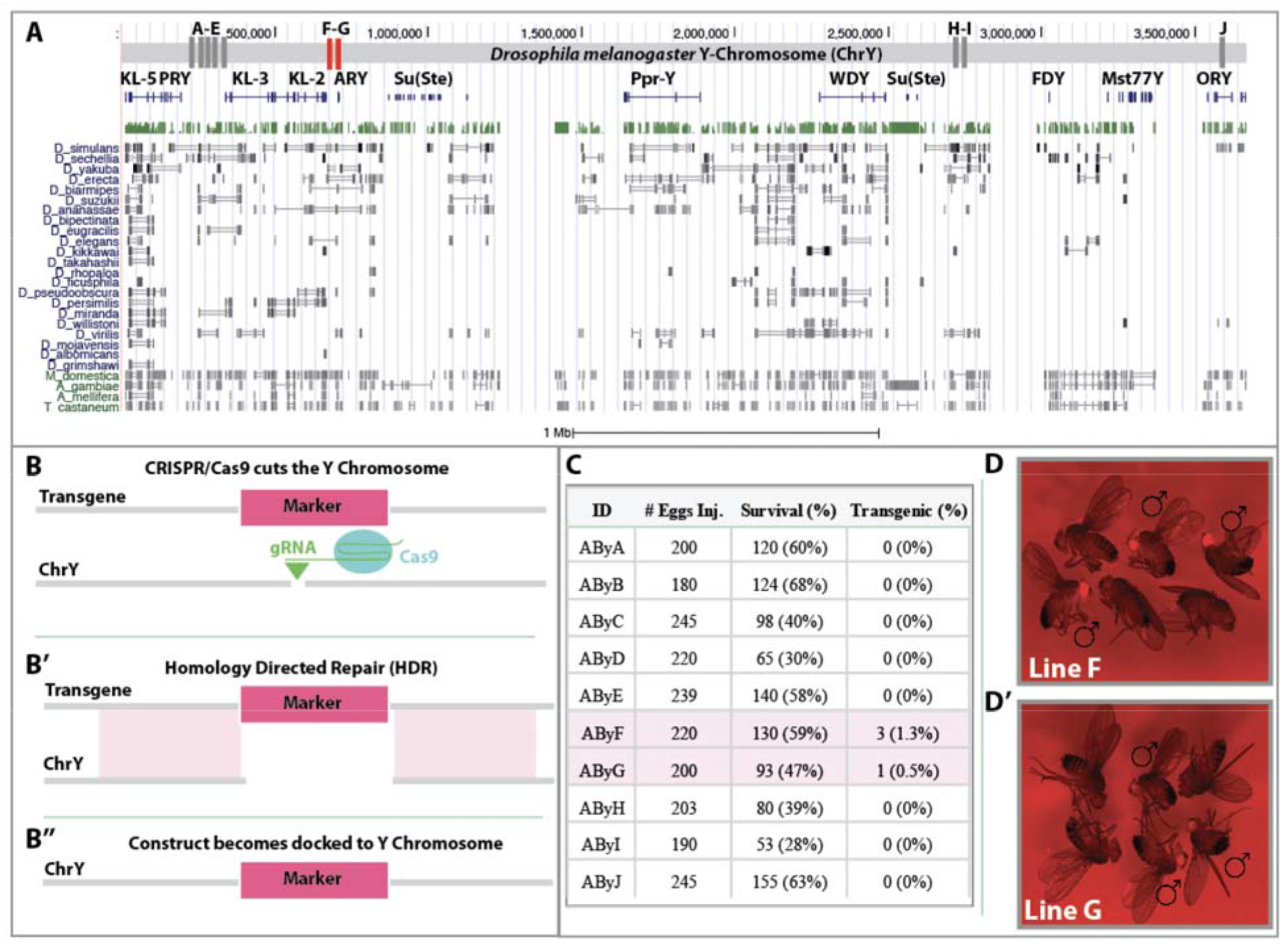
Targeted transgene insertion on the *D. melanogaster* Y-chromosome. (A) A map of the *D. melanogaster* Y-chromosome (adapted from the UCSC Genome Browser) shows the position of attempted transgene insertions (vertical bars labelled A-J; red bars indicate successful insertions, while grey bars indicate insertion failure) relative to the location of major genes. Conservation of gene regions in 26 other species is also indicated. (B) A schematic of the basic transgene utilized is shown. The transgene contains a marker flanked by homology arms to a specific region of the Y-chromosome, and is injected with a sgRNA that targets a sequence on the Y-chromosome between the homology arms, and a source of Cas9 (B). Following a Cas9-induced double-strand DNA break (DSB), the cell’s homology-directed repair (HDR) pathway utilizes the transgene as a repair template to copy the marker between the regions of homology (B’), generating a Y-chromosome with a marker gene at the precise sgRNA-targeted site (B’’). (C) The number of embryos injected for each of the Y-chromosome targeting transgenes, survival to larval stage of injected embryos, and transgenesis rate (# of independent transgenic individuals found/# of embryos injected) is shown. (D) All males (and none of the females) for each of the two recovered transgenic lines (AByF (D) and AByG (D’)) express eye-specific red fluorescence.

**Figure 2.**
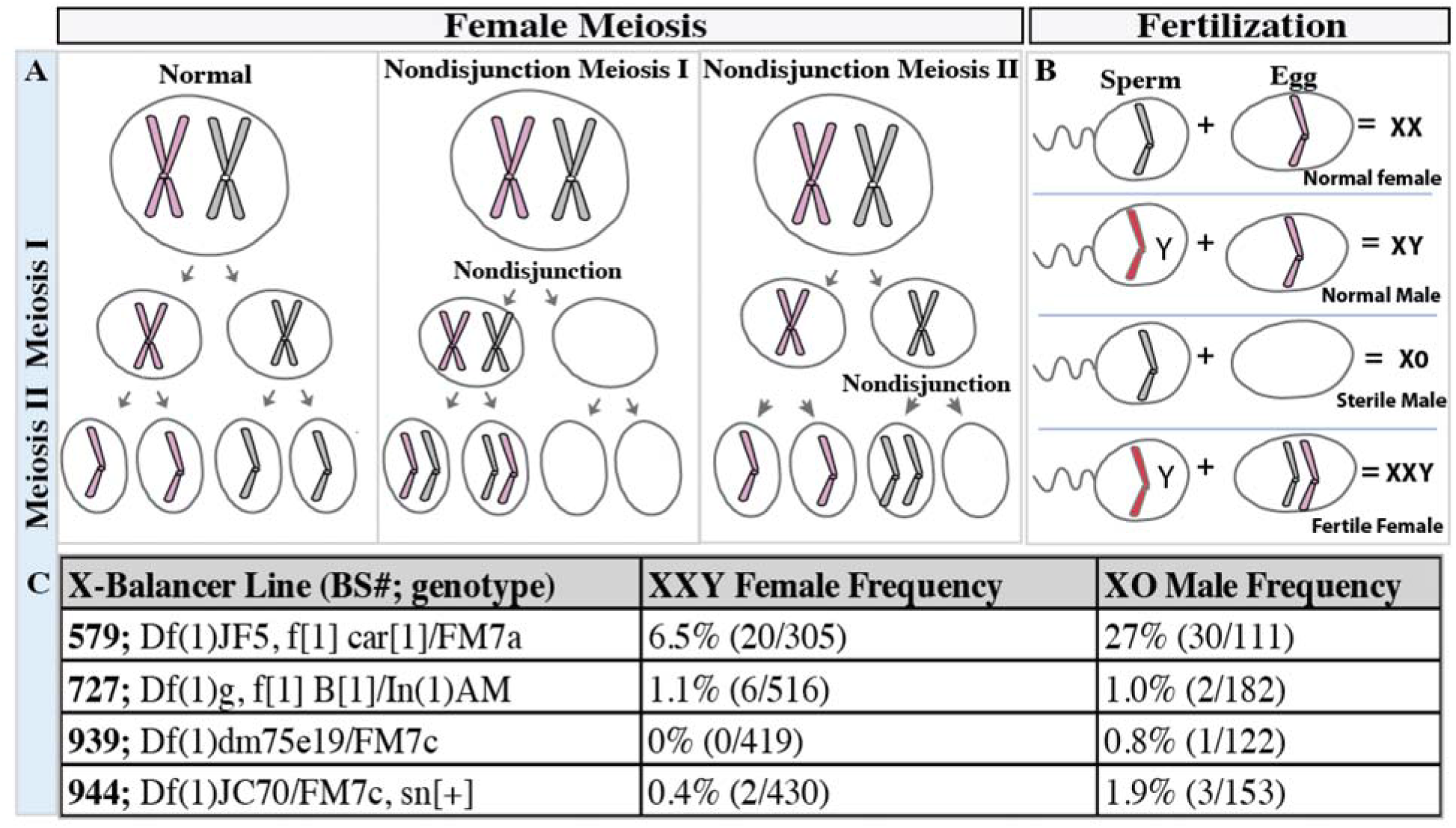
Using transgenic marked Y-chromosomes to track X-chromosome meiotic nondisjunction events. (A) In *D. melanogaster*, chromosome nondisjunction can occur in meiosis I or meiosis II; in either case, nondisjunction of X-chromosomes leads to formation of two types of aberrant gametes - ones that have an extra copy of the X-chromosome, and ones that lack an X-chromosome entirely. (B) If nondisjunction occurs during meiosis in a female, she will produce four types of viable offspring when outcrossed to a normal male: normal (XX) females, normal (XY) males, sterile (XO) males, and fertile (XXY) females. (C) Outcrosses of a marked Y-chromosome line to four distinct X-chromosome balancer lines (leftmost column) resulted in progeny arising from nondisjunction events. Frequencies of XXY females (females with marked Y-chromosome; middle column) and XO males (males with no Y-chromosome; rightmost column) are shown.

Additionally, the ability to express specific transgenes directly from the Y-chromosome would be useful in several different contexts. It would enable general studies of Y-chromosome gene function, for example, as well as analyses of chromosome loss and interaction (Szabad et al. 2012; Soós & Szabad 2014; Szabad & Würgler 1987). Perhaps more interestingly, it could aid in the development of genetic control mechanisms for insect vectors and pests, which offer promising solutions to significant health and agricultural problems (Champer et al. 2016; Esvelt et al. 2014; Sinkins & Gould 2006). For example, distortion of the sex ratio in favor of males can lead to a gradual population reduction and eventual elimination of a target population (Gould & Schliekelman 2004; Hamilton 1967; Hickey & Craig 1966; Papathanos et al. n.d.), and natural so-called meiotic driving Y-chromosomes have been described (Wood & Newton 1991; Newton et al. 1976; Sweeny & Barr 1978). A system for sex-ratio distortion can also be engineered by designing transgenes that target the X-chromosome during spermatogenesis, and such an X-shredder element has been developed in one species of mosquito (Windbichler et al. 2008; Galizi et al. 2014; Galizi et al. 2016). However, to be maximally effective and self-perpetuating, such an element would have to be linked to the Y-chromosome (Deredec et al. 2008; Deredec et al. 2011; Galizi et al. 2016), a feature that the developed systems lacked but that could be achieved by an efficient method of targeted Y-chromosome transgenesis.

Here we describe the development of a CRISPR/Cas9-based technique for site-specific engineering of the *D. melanogaster* Y-chromosome. Specifically, we demonstrate the ability to insert a fluorescently marked transgenic cassette at specific locations on the Y-chromosome, and to utilize the marked Y-chromosome to identify XXY females and XO males. The interspecies portability and limitations of this technique are also discussed.

## Results & Discussion

To generate transgenic elements that could be inserted in specific locations on the *D. melanogaster* Y-chromosome with CRISPR/Cas9-mediated HDR, we engineered a vector comprising a fluorescent marker (tdTomato) driven by the eye-specific 3xP3 promoter (Berghammer et al. 1999) and flanked by the *gypsy* (Holdridge & Dorsett 1991) and CTCF (Kyrchanova et al. 2008) insulators, with unique restriction sites upstream and downstream for cloning specific homology arms (Figure 1). We then selected ten distinct intergenic regions spanning the Y-chromosome for targeting (Figure 1A), identified a suitable sgRNA target site in each region, and cloned in homology arms, corresponding to ~800-1,000 base pairs of sequence 5’ and 3’ of each selected target site, upstream and downstream of the insulator-flanked 3xP3-tdTomato element to generate ten unique Y-chromosome targeting transgenes (AByA-J) (Figure 1A).

Each transgene was then injected, along with the appropriate *in vitro* transcribed sgRNA and Cas9 protein, into a transgenic line expressing a germline source of Cas9 using standard procedures (Gratz et al. 2014), and G_1_ progeny were screened for presence of eye-specific tdTomato fluorescence (Figure 1B). Of the ten distinct construct injections, only two (AByF and AByG) yielded transgenic male individuals for a success rate of 20%, despite the large number (>200 in most cases, 2142 total for all constructs) of G_0_ embryos injected for each construct (Figure 1C-D). This observation suggests that either the rate of Y-chromosome transgenesis is substantially lower than the rates reported in other studies utilizing CRISPR/Cas9-based HDR in autosomal locations (e.g., Gratz et al. 2014), or that a number of tested Y-chromosome target sites were not located in regions conducive to somatic gene expression. Neither of these explanations would be surprising, given the heterochromatic nature of the *D. melanogaster* Y-chromosome (Carvalho 2002), and simply suggests that several target regions must be tested in order to identify those suitable for HDR and/or that allow robust expression. Following their recovery, we outcrossed transgenic males to white eye (w-, w[1118]) females for several generations to confirm paternal inheritance of the tdTomato marker, and saw expected inheritance patterns (i.e., all male progeny and no female progeny of a male inherited the fluorescent marker). We also verified that the transgene was correctly inserted by performing PCR across the transgene-insertion junction on genomic DNA of transgenic males and sequencing the products.

We then set out to determine whether a marked Y-chromosome could be useful in identifying the rare progeny resulting from meiotic sex chromosome nondisjunction events in females (Figure 2A-B), as a clear means for identification of such individuals could be useful both for those that want to avoid them and those that wish to utilize them for addressing various research questions (e.g., Brown & Bachtrog 2017). To do this, we crossed males from the stronger-expressing of our two transgenic Y-chromosome lines, AByF, to a variety of X-chromosome balancer line virgins containing genetic elements such as inversions and rearrangements that are known to increase the probability of nondisjunction (Xiang & Hawley 2006; Gilliland et al. 2014). For four of these crosses, we identified progeny that appeared to be the result of meiotic nondisjunction events in the X-chromosome balancer females - XXY-bearing females (identified by presence of the fluorescent marker) and XO males (identified by the absence of the fluorescent marker) - at various frequencies ranging from 0-27% (Figure 2C). These aberrant progeny were individually outcrossed to a common laboratory stock (w[1118]) to test for fertility. All putative XO males were sterile (n=36) as expected (Griffiths 2005). All XXY females were fertile (n=28), and gave rise to progeny classes that indicated occurence of secondary nondisjunction in XXY parents, which is in agreement with previous observations (Bridges 1916; Xiang & Hawley 2006; Carpenter 1973). The presence or absence of the transgenic Y-chromosome in XXY females and XO males, respectively, was verified with PCR as described above. Given the high rates of nondisjunction individuals obtained by crossing the marked Y-chromosome line with certain X-chromosome balancers (e.g., balancer line 579), future experiments requiring large quantities of XXY females and XO males could be conducted using this approach, which could accelerate basic research on deciphering key biological processes and gene networks involved in Y-chromosome functionality and evolution.

In short, here we describe the engineering of two Y-chromosome marked transgenic *D. melanogaster* lines using site-specific, CRISPR/Cas9-mediated insertion. This may be useful for generating, identifying, and tracking progeny arising from rare meiotic nondisjunction events in a more straightforward manner than typically used (Brown & Bachtrog 2017; Piergentili 2010; Hess & Meyer 1968). More broadly, however, we provide a proof of principle example of a CRISPR/Cas-based strategy for docking transgenes site-specifically on the Y-chromosome that should in principle be applicable to many other species. Although the often-heterochromatic nature of a Y-chromosome may, as we observed here, reduce the number of targeted insertion sites that are amenable to modification and/or robust transgene expression, the ease with which distinct Y-chromosome targeting transgene cassettes can be generated should allow for testing of many possible insertion sites, greatly increasing the chances of successful transgene integration with desired expression. The ability to mark, and express transgenes directly from, the Y-chromosome not only allows for facile identification of Y-chromosome bearing individuals at various life stages and enables researchers to probe the enigmatic function of the Y-chromosome (e.g., (Brown & Bachtrog 2017; Bernardini et al. 2014), but could also help pave the way for engineering genetic insect vector and pest control strategies such as X-chromosome shredding (Hickey & Craig 1966; Hamilton 1967; Gould & Schliekelman 2004; Huang et al. 2007; Galizi et al. 2014; Champer et al. 2016; Galizi et al. 2016). This approach in particular depends on the destruction of X-bearing sperm to produce males that only give rise to male progeny (Huang et al. 2007; Champer et al. 2016), and requires the ability to meiotically express an X-chromosome targeting element from the Y-chromosome (Beaghton et al. 2016; Galizi et al. 2016). The strategy described here could be used to dock such an X-chromosome targeting transgene (perhaps one based on CRISPR/Cas9 technology, Galizi et al. 2016; Zuo et al. 2017) to the Y-chromosome of many insects, including the malaria vector *Anopheles gambiae*, to facilitate the engineering of the promising X-chromosome shredding genetic control strategy that can be used to combat malaria and other vectored pathogens (Gould & Schliekelman 2004; Hall et al. 2016; Papathanos et al. n.d.; Champer et al. 2016).

## Experimental Procedures

### Construct Design and Assembly

To generate vector ABy, the base vector used to generate vectors AByA-AByJ, several components were cloned into the *piggyBac* plasmid pBac[3xP3-DsRed] (Li, Bui, Yang, et al. 2017) using Gibson assembly/EA cloning (Gibson et al. 2009). First, a Gypsy insulator fragment amplified with primers ABy.1 and ABy.2 from *Drosophila* genomic DNA, the 3xP3 promoter amplified with primers ABy.3 and ABy.4 from plasmid pBac[3xP3-EGFP afm] (Horn & Wimmer 2000), and a *Drosophila* codon optimized tdTomato marker amplified with primers ABy.5 and ABy.6 from a gene synthesized vector (Genscript, Piscataway, NJ) were cloned into a BstBI/NotI digested pBac[3xP3-DsRed] backbone using EA cloning. The resulting plasmid was digested with AvrII, and the following components were cloned in via EA cloning: an *attP* sequence from plasmid M{3xP3-RFP attP}(Bischof et al. 2007) amplified with primers ABy1.7 and ABy.8, and a CTCF insulator fragment amplified with primers ABy.9 and ABy.10 from *Drosophila* genomic DNA. All primer sequences are listed in Supplementary Table 1.To generate the final vectors containing homology arms specific to each putative Y-chromosome docking site, vector ABy was first digested with PmeI and each 5’ homology arm (amplified with primers specific to the Y docking site from *Drosophila* genomic DNA) was individually cloned in using EA cloning. Each resulting intermediate plasmid was then digested with EcoRI, and each corresponding 3 ‘ homology arm (amplified with primers specific to the Y docking site from *Drosophila* genomic DNA) was cloned in using EA cloning. The 10 Y-chromosome docking site-specific construct names and the primers used to amplify the 5’ and 3’ homology arms for each are listed in Supplementary Table 1. Vectors AByG and AByH are available from Addgene (#s 111084 and 111083, respectively). To generate double-stranded DNA breaks for vector incorporation, sgRNAs targeting 10 distinct regions of the Y-chromosome were *in vitro* transcribed with the MEGAscript™ T7 Transcription Kit (ThermoFisher Scientific, cat. # AM1334) using a self-annealing set of primers, with a unique forward primer for each target site and a universal reverse primer. Genomic locations of each sgRNA target sequence and primers used for *in vitro* transcription are listed in Supplementary Table 1.

### Fly Culture and Strains

Fly husbandry and crosses were performed under standard conditions at 25°C. Rainbow Transgenics (Camarillo, CA) carried out all of the fly injections. Each construct was pre-mixed with Cas9 protein (PNA Bio Inc., cat. # CP01-20) and *in vitro* transcribed sgRNAs at the following concentrations: 400ng/μL plasmid, 40ng/μL sgRNA, and 300ng/μL Cas9 protein. Constructs were injected into a vasa-Cas9 transgenic line marked with 3xP3-GFP (Bloomington Drosophila Stock Center #51324, w[1118]; PBac{y[+mDint2]=vas-Cas9}VK00027). Only transgenic males were recovered, and these were singly outcrossed to w-(w[1118]) virgin females to verify paternal transmission. Male progeny without the vasa-Cas9 transgene were selected and further crossed to w-to establish a stock. Stocks for the two transgenic lines recovered (AByF and AByG) were submitted to the Bloomington Drosophila Stock Center.

### Molecular characterization of Y-chromosome lines

To confirm correct insertion of transgenes on the Y-chromosome, PCRs were carried out on genomic DNA from transgenic males (Figure 1). Briefly, genomic DNA was extracted from individual flies with the DNeasy Blood & Tissue kit (QIAGEN, cat. # 69504) following the manufacturer’s protocol. PCR was carried out using standard procedures to amplify the junction of the transgenes and the surrounding genomic region (as well as the unmodified target genomic region as a negative control and a region of the X-chromosome as a positive control) using primers listed in Supplementary Table 1. The PCR program utilized was as follows: 98°C for 30 seconds; 35 cycles of 98°C for 10 seconds, 56°C for 20 seconds, and 72°C for 30 seconds; then 72°C for 10 minutes. PCR products were purified with the MinElute PCR Purification Kit (QIAGEN, cat. #28004) according to the manufacturer’s protocol and sequenced with Sanger sequencing (SourceBioScience) utilizing the same primers as used for the PCRs. Sequences were analyzed with DNAStar software.

### Generation and characterization of XXY females and XO males

To determine whether a marked Y-chromosome could be utilized to identify individuals that result from meiotic nondisjunction events, a single male from the AByF transgenic line (which showed stronger fluorescent marker expression levels than line AByG transgenic males) was crossed to virgins from the following X-chromosome balancer stocks from the Bloomington Drosophila Stock Center (BSC): BSC #s 579, 723, 727, 939, 944, 946, 959, 976, 3347, 6002, 6007, 6019, 6219, 7200, and 27887. Each cross was set in triplicate. Progeny were screened to identify females with fluorescent red eyes (putative XXY individuals) and males without fluorescent red eyes (putative XO individuals). Frequency of putative XXY and XO individuals was calculated by dividing the total number of each individual type found for each type of cross by the total number of females or males, respectively (Figure 2C). Each XO individual was outcrossed to w- (w[1118]) virgins to test for fertility. To molecularly confirm the presence of the transgenic Y-chromosome in putative XXY females and the absence of any Y-chromosome in putative XO males, genomic DNA was extracted as above, and PCRs were performed as above with primers utilized for the AByF 5’ gDNA-transgene junction, the AByF 5’ gDNA-transgene junction, the AByF wildtype locus, and the X-chromosome positive control locus (Supplementary Table 1).

## Acknowledgements

This work was supported in part by funding from the California Cherry board, an NIH-K22 Career Transition award (5K22AI113060), an NIH Exploratory/Developmental Research Grant Award (1R21AI123937), and a Defense Advanced Research Project Agency (DARPA) Safe Genes Program Grant (HR0011-17-2-0047) awarded to O.S.A. Stocks obtained from the Bloomington Drosophila Stock Center (NIH P40OD018537) were used in this study. We thank Thom Kaufman (Indiana University Bloomington, Indiana) for useful discussions and for providing the BSC X-chromosome balancer lines.

## Author Contributions

O.S.A and A.B. conceived and designed experiments. A.B. performed molecular and genetic experiments. All authors contributed to the writing, analyzed the data, and approved the final manuscript.

